# Adverse childhood experiences in families with multiple members diagnosed to have psychiatric illnesses

**DOI:** 10.1101/745521

**Authors:** Amala Someshwar, Bharath Holla, Preeti Pansari Agarwal, Anza Thomas, Anand Jose, Bobin Joseph, Birudu Raju, Hariprasad Karle, M Muthukumaran, Prabhath G Kodancha, Pramod Kumar, Preethi V Reddy, Ravi Kumar Nadella, Sanjay T Naik, Sayantanava Mitra, Sreenivasulu Mallappagiri, Vanteemar S Sreeraj, Srinivas Balachander, Suhas Ganesh, Pratima Murthy, Vivek Benegal, Janardhan Y. C. Reddy, Sanjeev Jain, Jayant Mahadevan, Biju Viswanath

**Affiliations:** Department of Psychiatry, National Institute of Mental Health and Neuro Sciences (NIMHANS), Bengaluru, India; The National Centre for Biological Sciences (NCBS), Bengaluru, India; National Institute of Mental Health and Neuro Sciences (NIMHANS); Institute for Stem Cell Biology and Regenerative Medicine (InStem); National Center for Biological Sciences (NCBS)

**Author notes:** Corresponding author Dr. Jayant Mahadevan, Assistant Professor of Psychiatry, National Institute of Mental Health and Neurosciences, Bangalore, India 560029, Dr. Biju Viswanath, Associate Professor of Psychiatry, National Institute of Mental Health and Neurosciences, Bangalore, India 560029. Equal contribution (Amala Someshwar, Bharath Holla).

**Keywords:** adverse childhood experiences (ACE), psychiatric disorder, multiplex families, early onset, gender differences

## Abstract

**Objective:** Adverse Childhood Experiences (ACEs) are linked to the development of a number of psychiatric illnesses in adulthood. Our study examined the pattern of ACEs and their relation to the age of onset (AAO) of major psychiatric conditions in individuals from families that had ≥ 2 first degree relatives with major psychiatric conditions (multiplex families) identified as part of an ongoing longitudinal study.

**Methods:** Our sample consisted of 509 individuals from 215 families. Of these, 268 were affected i.e diagnosed with bipolar disorder (BPAD) (*n*=61), obsessive-compulsive disorder (OCD) *(n*=58), schizophrenia (*n*=52), substance dependence (SUD) (*n*=59), or co-occurring diagnoses (*n*=38); while 241 were at-risk first degree relatives (FDRs) who were either unaffected (*n*=210) or had other depressive or anxiety disorders (*n*=31). All individuals were evaluated using the Adverse Childhood Experiences – International Questionnaire (ACE-IQ) and ACE binary and frequency scores were calculated.

**Results:** It was seen that affected males, as a group, had the greatest ACE scores in our sample. A cox mixed-effects model fit by gender revealed that higher ACE binary and frequency scores were associated with significantly increased risk for an earlier AAO of psychiatric diagnoses in males. A similar model that evaluated the effect of diagnosis revealed an earlier AAO in OCD and SUD, but not in schizophrenia and BPAD.

**Conclusions:** Our study indicates that ACEs brought forward the onset of major psychiatric conditions in men and in individuals diagnosed with OCD and SUD. Ongoing longitudinal assessments in FDRs from these families are expected to identify mechanisms underlying this relationship.

## Introduction

Adverse childhood experiences (ACEs) occur often (Kessler et al., 2010; Soares et al., 2016), and have been linked to a broad range of negative outcomes, both in terms of mental (Edwards et al., 2003; Van der Kolk, 2017) and physical health (Van der Kolk, 2017; Anda et al., 2006; Perry et al., 1995), as well as quality of life and life experiences (Chapman et al., 2011; Hillis et al., 2004). Overall, with respect to mental health, individuals who reported being physically and sexually abused as children were found to have more psychiatric conditions as adults (Jasinski et al., 2000; Leverich et al., 2002). Women who reported being victims of childhood sexual assault were found later to report greater levels of anxiety, anger regulation, paranoid ideation, and obsessive-compulsive symptoms (Murphy et al., 1988).

In terms of specific psychiatric disorders, a number of ACEs have been associated with the severity of specific disorders such as addiction, major depression, and obsessive compulsive disorder (Brodsky et al., 2001; Dube et al., 2003; Lochner et al., 2002). The association between ACEs and the onset of bipolar disorder and schizophrenia has also been previously investigated, with presence of ACEs being associated with an increased risk of psychosis (Etain et al., 2013; Read et al., 2005; Watson et al., 2014). Exposure to certain types of ACEs as well as ACEs at a greater intensity may be related to an earlier onset age of psychiatric conditions, such as substance dependence, depression, and schizophrenia (Etain et al., 2013; Read et al., 2005; Bernet and Stein, 1999; Li et al., 2012), which may, in turn, indicate a more severe phenotype (Kessler et al., 2007). Early exposure to trauma also appears to increase risk of psychotic symptoms in at-risk adolescents (Spauwen et al., 2006).

Overall, childhood trauma seems to contribute to the future occurrence of diverse symptom clusters, and possibly to an earlier age of onset of illness. In this study, we examine the effect of ACEs on age of onset of different psychiatric syndromes. A trans-diagnostic approach could help us to understand the differential effect of ACEs on the age of onset of such syndromes. In addition, evaluating individuals from multiplex families with a pre-existing genetic loading for psychiatric illness (see under ‘methods’ the source of data), might inform us about the role of ACEs in bringing forward the onset of illness in already vulnerable individuals.

The current study describes the pattern of ACEs in affected individuals and at-risk first degree relatives (FDRs) from multiplex families with major psychiatric disorders [schizophrenia, bipolar disorder (BPAD), substance dependence (SUD), obsessive compulsive disorder (OCD)] and its relationship to age of onset of the disorders. We hypothesized that ACEs would influence the age of onset of different psychiatric syndromes in individuals from multiplex families.

## Methods

### Procedure

This study draws on data from the Accelerator program for Discovery in Brain disorders using Stem cells (ADBS) project, an ongoing longitudinal study at the National Institute of Mental Health and Neurosciences (NIMHANS), the National Centre for Biological Sciences (NCBS) and the Institute for Stem Cell Science and Regenerative Medicine (InStem), that began in 2016 (Viswanath et al., 2018). The project was reviewed and approved by the institutional ethics review board and written informed consent was obtained from all individuals who were recruited.

This study includes families in whom multiple members (at least 2 affected first degree relatives in a nuclear family) are diagnosed to have a major psychiatric disorder (schizophrenia, BPAD, OCD, Alzheimer’s dementia and SUD). The study comprised of two levels of assessment – a brief assessment, and the neurodevelopmental endo-phenotype (deep) assessment. Overall health assessments were also performed on all participants, in order to take note of any pre-existing medical conditions. All psychiatric diagnoses were corroborated by two trained psychiatrists using the Mini International Neuropsychiatric Interview (MINI) (Sheehan et al., 1998). Further clinical evaluation, including assessment of temperament, personality, adverse childhood experiences, life events, handedness, functioning, and psychopathology specific scales, were done on participants who consented to the deep assessments. For the present analysis, the adverse childhood experiences international questionnaire (ACE-IQ) (World Health Organization, 2011), and basic sociodemographic and clinical information (including onset age of the full syndrome, gender, and maximum education attained, and psychiatric diagnosis) were used. Individuals affected with Alzheimer’s dementia were excluded from the study, as most were unable to complete the ACE-IQ form. A total of 509 participants (affected individuals and at-risk FDRs) met these criteria from the data available in the ADBS project. For the purpose of the present study, we divided the affected individuals into five groups by diagnosis: BPAD, OCD, schizophrenia, SUD, and co-occurring diagnoses group (American Psychiatric Association, 2000). Co-occurring diagnoses were defined as presence of more than one lifetime Axis I diagnosis of either BPAD, SUD, OCD, or schizophrenia. At-risk FDRs consisted of individuals with no Axis I diagnoses, as well as individuals with Axis I diagnoses other than the diagnostic groups described above.

### Measures

*Age of Onset (AAO)* was defined as the age at which individuals fulfilled DSM IV TR criteria for any of the four disorders, as determined by the psychiatrist, based on the information obtained from the patient and their family members.

*Adverse Childhood Experiences* were assessed using the ACE-IQ questionnaire. It consists of 31 questions on 13 subdomains: physical abuse; emotional abuse; contact sexual abuse; alcohol and/or drug abuser in the household; incarcerated household member; household member with a psychiatric condition or suicidality; household member treated violently; one or no parents, parental separation or divorce; emotional neglect; physical neglect; bullying; community violence; and collective violence (World Health Organization, 2011).

Binary ACE Score was calculated as the total number of subdomains where adversity was reported, independent of frequency. Frequency ACE Score was calculated as the total number of subdomains where adversity was reported at a predefined frequency, as specified in the WHO ACE-IQ guidelines (World Health Organization, 2011). Both binary and frequency ACE scores ranged from 0 to 13. The frequency version of the score represents more severe form of the childhood adversity.

### Statistical Analyses

Means, medians and proportions were used to describe the sociodemographic details and ACE binary and frequency scores of the study sample. Generalized linear mixed effects Poisson regression was used to examine group-by-gender interaction for ACE binary and frequency scores, after accounting for correlated ACEs between family members. We applied mixed effects survival analysis methodology to model AAO, using correlated frailty models to account for correlated AAO between family members. Here, survival analyses models AAO as a function of time, and takes into account that some records are uncensored reflecting the actual AAO (affected individuals) and some are censored at the age last seen as unaffected (at risk FDRs). Separate cox mixed effects models were fit to examine the effects of both ACE Scores, ACE-by-Gender interactions, ACE-by-diagnosis interactions on the AAO of psychiatric illness. The hazard ratios (HR) provides the instantaneous risk of becoming affected at age *‘t’*, given that an individual was not affected at age (t-1) as an effect of increase in ACE scores. In each case, the models were also repeated after excluding individuals with AAO <18 years.

All analyses were performed in R environment for statistical computing, version 3.5.2, using the packages ‘lme4’ and ‘coxme’.

## Results

### Sociodemographic and Clinical Details

This study sample consisted of 509 individuals (43.2% females) from 215 different families. Of the participants, 268 were affected (36.6% females), while 241 were at-risk FDRs (50.6% females). *Supplementary Table 1*, describes the sociodemographic characteristics of the individuals in the sample. The mean AAO for the whole sample was 31.8 ± 14.1 years. The mean AAO for men was 29.8 ± 13.9 years and for women was 34.4 ± 14.0 years. *Supplementary Table 2*, describes clinical details of the sample, by diagnosis.

### Description of ACE

A total of 450 out of 509 (88.4%) individuals reported having experienced at least one ACE, indicated by an ACE binary score of ≥1; 404 out of 509 (79.4%) experienced at least one ACE frequently, indicated by an ACE frequency score of ≥1 (*Figure 1A and 1B*). The mean (*SD*) ACE binary scores for individuals with BPAD, schizophrenia, OCD, SUD and co-occurring diagnoses were 3.2 (2.1), 3.13 (2.5), 2.98 (1.8), 5.57 (2.8), and 4.02 (2.4) respectively. The mean (*SD*) ACE frequency scores for individuals with BPAD, schizophrenia, OCD, SUD and co-occurring diagnoses were 1.9 (1.6), 2.0 (2), 1.71 (1.4), 3.8 (2.5), and 2.73 (1.8) respectively. *Figure 1C and 1D* describe the distribution of ACE binary and frequency scores in the affected group split by diagnosis.

**Figure 1:**
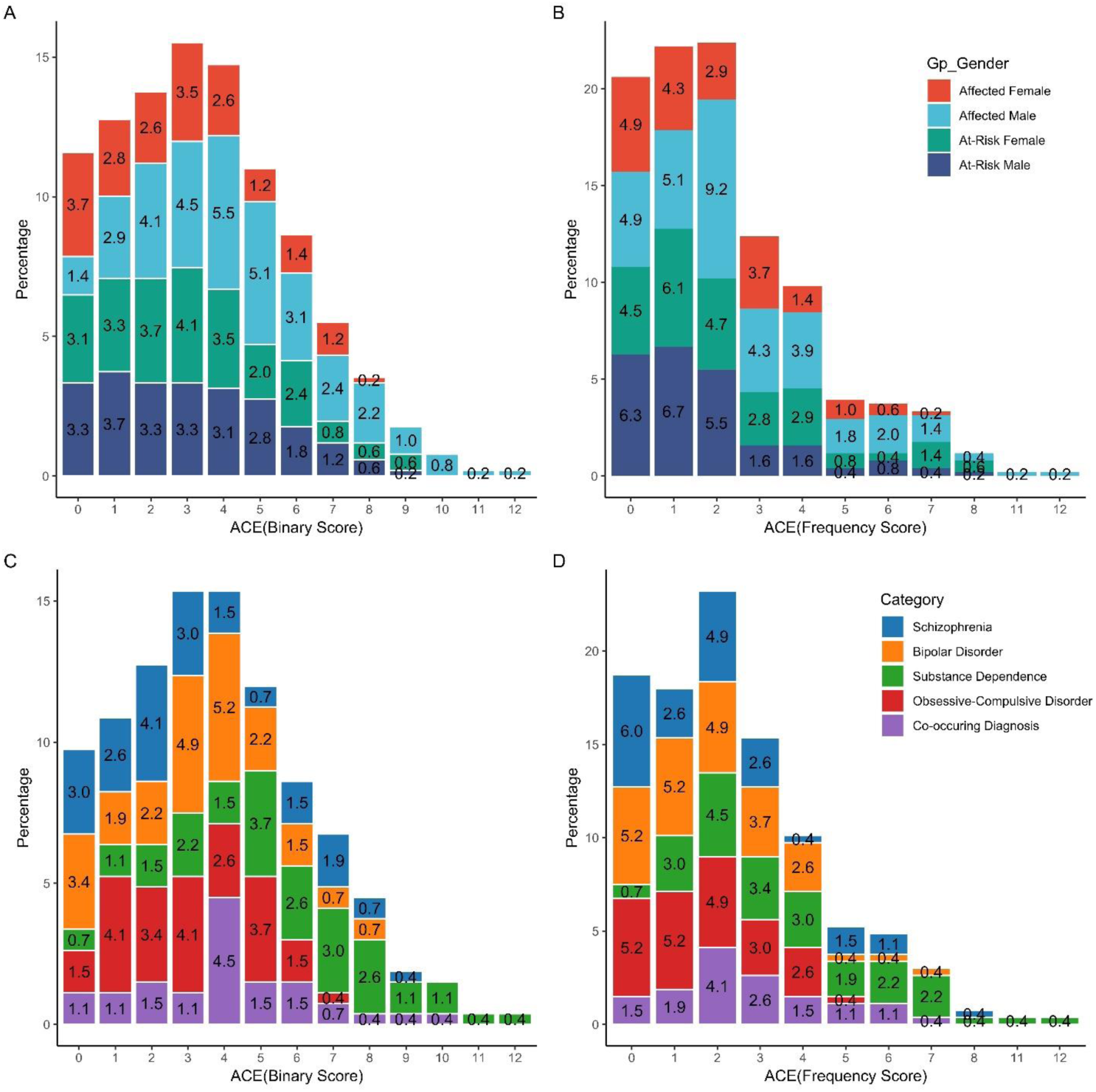
Distribution of ACE Binary and Frequency Scores for (A-B) affected individuals and at risk first-degree relatives split by gender and (C-D) affected individuals split by diagnosis.

*Figure 2A and 2B* describe the incidence rate (95%CI) of ACE binary and frequency scores in affected and at-risk groups split by gender. Affected males had significantly greater ACE binary and frequency scores compared to affected females, at-risk males or females.

**Figure 2:**
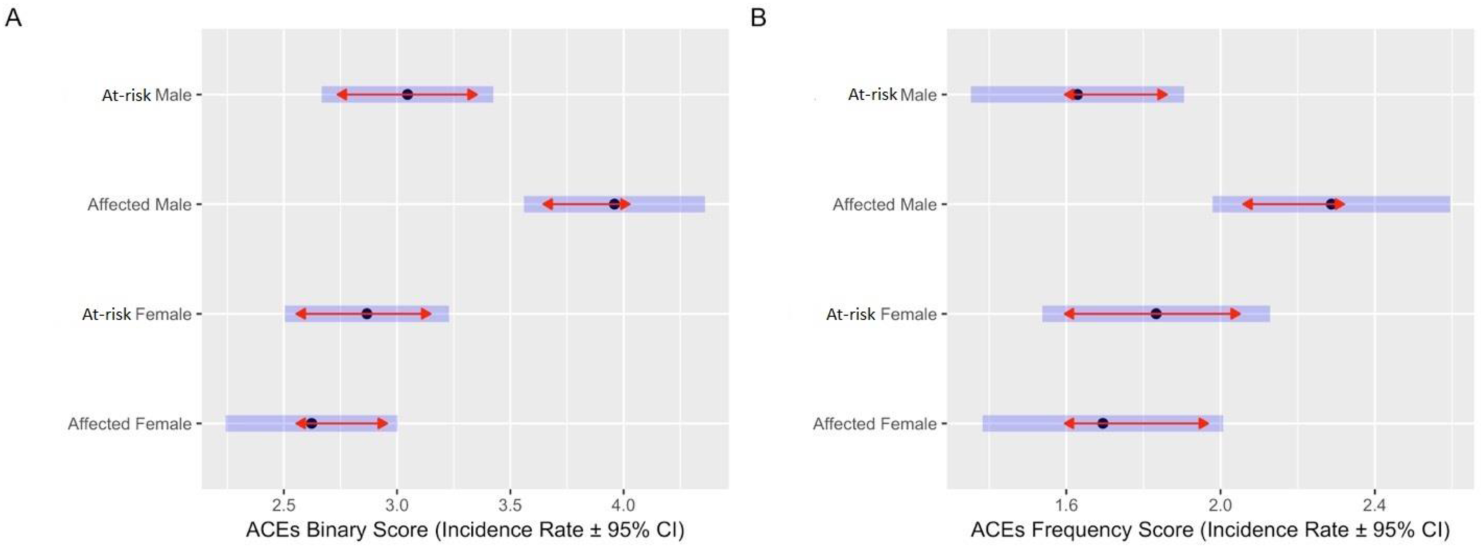
Incidence rate ± 95% Confidence Interval (CI) for the Group-by-Gender interaction for (A) ACE Binary and (B) ACE Frequency Scores. The results indicate significantly greater incidence rates of both Binary and Frequency ACE scores for Affected Males. Legend: Incidence rate ± 95% Confidence Interval (CI) are back-transformed from the log scale and represented by dot and blue bars. The red arrows are for the post-hoc tukey comparisons, with overlaps in the arrow indicating that the between-group difference for that comparisons is not significant.

### Association between AAO and ACE Scores *(Table 1)*

Models 1 and 2 investigated the effects of ACEs on the age of onset of psychiatric illnesses and found that greater ACE binary (HR = 1.118, 95% CI = 1.066 - 1.172) and frequency (HR = 1.103, 95% CI = 1.044 - 1.165) scores significantly increased the risk for an earlier age of onset across all disorders. As the ACE-IQ form pertains to incidents occurring below the age of 18, we also repeated the analysis after excluding participants with an onset age ≤ 18. This was to adjust for the influence of ACEs that may have occurred as a consequence of the illness. The associations remained significant for the binary (HR = 1.131, 95% CI = 1.068 - 1.197) and frequency (HR = 1.126, 95% CI = 1.055 - 1.201) scores.

**Table 1:** Results from Mixed Effects Cox Models (1-4) to examine the relationship between ACE Scores, ACE-by-Gender interactions on the Age-at-Onset (AAO) of psychiatric illness. The unaffected individual from the high-risk families are censored at the age last seen. Family ID was used as random effects confounding variable. Models (1-4) were repeated after excluding individuals with AAO <18 years, to adjust for the influence of ACEs that may have occurred as a consequence of the illness.

|  |  | Full Sample (events, n= 268, 509) |  |  | Sample with AAO >18y (events, n = 192, 433) |  |  |
| --- | --- | --- | --- | --- | --- | --- | --- |
| No | Model predictors | HR (95% CI) | z | p.value | HR (95% CI) | z | p.value |
| ACE |  |  |  |  |  |  |  |
| 1 | ACE(Bin) | 1.118 (1.066 - 1.172) | 4.57 | <0.001 | 1.131 (1.068 - 1.197) | 4.23 | <0.001 |
| 2 | ACE(Freq) | 1.103 (1.044 - 1.165) | 3.49 | <0.001 | 1.126 (1.055 - 1.201) | 3.57 | <0.001 |
| ACE-by-Gender interaction |  |  |  |  |  |  |  |
| 3 | ACE(Bin):Female | 0.996 (0.926 - 1.072) | -0.09 | 0.92 | 1.019 (0.937 - 1.107) | 0.44 | 0.66 |
|  | ACE(Bin):Male | 1.155 (1.102 - 1.211) | 6.00 | <0.001 | 1.175 (1.11 - 1.244) | 5.53 | <0.001 |
| 4 | ACE(Freq):Female | 0.954 (0.874 - 1.042) | -1.05 | 0.29 | 0.983 (0.891 - 1.086) | -0.33 | 0.74 |
|  | ACE(Freq):Male | 1.185 (1.121 - 1.252) | 5.97 | <0.001 | 1.226 (1.146 - 1.312) | 5.92 | <0.001 |

Bin, Binary and Freq, Frequency.

### Association between AAO and ACE by Gender Interactions *(Table 1)*

Models 3 and 4 investigated the effects of ACE-by-gender interactions on age of onset of psychiatric illness and found that greater ACE binary (HR = 1.155, 95% CI = 1.102 - 1.211) and frequency (HR = 1.185, 95% CI = 1.121 - 1.252) scores significantly increased the risk for an earlier age of onset only in males and not in females. The same held true even on excluding individuals with onset age ≤ 18. As there was a clear gender skew in individuals with substance dependence, this analysis was redone after excluding the substance dependence group; the gender difference continued to persist for both binary (HR = 1.132, 95% CI = 1.047 – 1.224, p = 0.002) and frequency (HR = 1.189, 95% CI = 1.072 – 1.319, p = 0.001) scores in males.

### Association between AAO and ACE by Diagnosis Interactions *(Table 2)*

Models 5 and 6 investigated the effects of ACE-by-diagnosis interactions on age of onset of psychiatric illness. It was found that greater ACE binary (HR = 1.254, 95% CI = 1.133 - 1.388) and frequency (HR = 1.336, 95% CI = 1.15 - 1.552) scores significantly increased the risk for an earlier age of onset in individuals with OCD both in the full sample and after exclusion of individuals with onset age ≤ 18. This finding was also seen for both ACE binary (HR = 1.123, 95% CI = 1.03 - 1.224) and frequency (HR = 1.143, 95% CI = 1.015 - 1.288) scores in individuals with co-occurring diagnoses but only in the full sample. In individuals with SUD, greater ACE binary (HR = 1.088, 95% CI = 1.015 - 1.166) and frequency (HR = 1.135, 95% CI = 1.038 - 1.24) scores predicted earlier age of onset in only in those with an onset age > 18 years and not in the full sample.

**Table 2:** Results from Mixed Effects Cox Model (5-6) to examine the relationship between ACE-by-Diagnosis interaction on the Age-at-Onset (AAO) of psychiatric illness. Family ID was used as random effects confounding variable. Models (5-6) were repeated after excluding individuals with AAO <18 years, to adjust for the influence of ACEs that may have occurred as a consequence of the illness. ACEs (Binary and Frequency) significantly increased the hazard ratio (HR) (21-34%) for an earlier AAO for both >18y and full sample for OCD. However, for SUD the increased HR were only seen in >18y sample and not the full sample, while for co-occurring diagnoses the increased HR was seen only for the full sample and not for >18y sample

| No | Model predictors | Full Sample (events, n = 268) |  |  | Sample with AAO >18y (events, n = 192) |  |  |
| --- | --- | --- | --- | --- | --- | --- | --- |
|  |  | HR (95% CI) | z | p.value | HR (95% CI) | z | p.value |
| 5 | ACE(Bin):SCZ | 1.032 (0.942 - 1.13) | 0.67 | 0.5 | 1.06 (0.954 - 1.178) | 1.08 | 0.28 |
|  | ACE(Bin):BPAD | 1.052 (0.962 - 1.151) | 1.11 | 0.27 | 0.927 (0.82 - 1.048) | -1.21 | 0.23 |
|  | ACE(Bin):SUD | 1.048 (0.989 - 1.112) | 1.58 | 0.11 | **1.088 (1.015 - 1.166)** | **2.4** | **0.017** |
|  | ACE(Bin):OCD | **1.254 (1.133 - 1.388)** | **4.39** | **<0.001** | **1.21 (1.048 - 1.397)** | **2.6** | **0.009** |
|  | ACE(Bin):Co-occ | **1.123 (1.03 - 1.224)** | **2.63** | **0.009** | 1.108 (0.988 - 1.242) | 1.76 | 0.078 |
| 6 | ACE(Freq):SCZ | 1.029 (0.91 - 1.163) | 0.45 | 0.65 | 1.091 (0.952 - 1.251) | 1.25 | 0.21 |
|  | ACE(Freq):BPAD | 1.044 (0.917 - 1.19) | 0.65 | 0.51 | 0.919 (0.774 - 1.091) | -0.97 | 0.33 |
|  | ACE(Freq):SUD | 1.059 (0.981 - 1.143) | 1.47 | 0.14 | **1.135 (1.038 - 1.24)** | **2.8** | **0.005** |
|  | ACE(Freq):OCD | **1.336 (1.15 - 1.552)** | **3.79** | **<0.001** | **1.315 (1.066 - 1.622)** | **2.55** | **0.011** |
|  | ACE(Freq):Co-occ | **1.143 (1.015 - 1.288)** | **2.2** | **0.028** | 1.146 (0.982 - 1.337) | 1.73 | 0.084 |

SCZ, schizophrenia; BPAD, bipolar affective disorder; SUD, substance use disorder; OCD, obsessive compulsive disorder; Co-occ, Co-occurring Diagnosis; Bin, Binary and Freq, Frequency.

## Discussion

The pattern of ACEs in affected and at-risk FDRs from multiplex families, and across diagnostic groups was evaluated. Further, we investigated the relationship between ACEs and the age of onset of psychiatric conditions.

We observed that in multiplex families, ACE binary and frequency scores were higher in affected individuals as compared to their at-risk FDRs. Specifically, they were highest for affected males in the sample. It was also seen that greater adversity scores increased the risk of an earlier AAO of psychiatric conditions by 11.8% (ACE binary) and 10.3% (ACE frequency), which was most evident in males [ACE binary: 15.5% and ACE frequency: 18.5%]. In terms of specific diagnosis, greater adversity scores increased the risk for an earlier age of onset in individuals with OCD [ACE binary: 25.4% and ACE frequency: 33.6%], co-occurring diagnoses [ACE binary: 12.3% and ACE frequency: 14.3%] and substance dependence (when the age of onset of dependence was >18 years) [ACE binary: 8.8% and ACE frequency: 13.5%].

ACE binary and frequency scores were higher for affected individuals than for at-risk individuals, and this observation is in line with previous studies (Leverich et al., 2002; Felitti et al., 2019; Dube et al., 2005). Since both affected and at-risk individuals in our study would have shared similar family environments, this difference could also be attributable to a greater recall of ACEs by affected individuals. It may also have been due to the fact that individuals who develop psychiatric illness in adulthood may manifest pre-clinical symptoms in childhood and adolescence, which expose them to a greater risk of experiencing adversity (Howes and Murray, 2014).

It was seen that higher ACE binary and frequency scores predicted an earlier age of onset across disorders. This illustrates that both the presence and the severity of adverse childhood experiences contribute to an earlier than usual age of onset. These findings are consistent with previous research which have found that the number and severity of childhood adverse events are a risk factor for onset of mental illness in adulthood (Edwards et al., 2003; Felitti et al., 2019; Bernet and Stein, 1999; van der Kolk, 2003).

Specifically, ACEs were seen to predict an earlier age of onset in individuals with OCD. Literature on the relationship between ACE and OCD is scarce and inconclusive (Selvi et al., 2012). Some studies report no association between ACE and OCD and few others support higher ACE in those with OCD, but possibly mediated by coexisting affective, anxiety, substance use and eating disorders(Visser et al., 2014; Briggs and Price, 2009; Benedetti et al., 2014). It is in this context, that an association between ACE and onset of OCD gains significance. Among these conditions, greater adverse events were found to predispose to an earlier age of onset for individuals with OCD. The association survived even after controlling for early onset OCD, implying the effect of ACE on onset of OCD irrespective of AAO. Our finding is also consistent with studies where individuals with OCD experience greater adversity than first degree relatives (Bey et al., 2017) and also have greater severity of future symptoms (Lin et al., 2007). It is possible that ACEs bring forward the onset of OCD in genetically vulnerable individuals.

ACEs were also seen to predict earlier age of onset of SUD, after excluding individuals in whom the age of onset was prior to 18 years. This confirms the observations made by previous studies in SUD (Anda et al., 2006; Dube et al., 2003; Dube et al., 2006), which also show a clear relationship between adversity and development of SUD. Anxiety and obsessionality, as well as addiction, are also known to have a complex pattern for clustering in families, with both genetic and non-genetic and learning factors being implicated (Mataix-Cols et al., 2013; Craske et al., 2017; Merikangas et al., 1998). ACEs also predicted earlier age of onset in individuals with co-occurring diagnoses, but this did not persist after exclusion of individuals with age of onset prior to 18 years, indicating that greater ACEs may have been a consequence, rather than antecedent, of the illness.

ACEs did not influence age of onset in schizophrenia and BPAD in our sample. This may be related to the fact that heritability estimates in schizophrenia (Hilker et al., 2018) and BPAD (McGuffin et al., 2003) are greater than those for OCD (Browne et al., 2014) and SUD (Ducci and Goldman, 2012). Since, individuals in our sample already had a strong genetic predilection the role of childhood environmental exposures such as ACEs may have been less significant for individuals with schizophrenia and BPAD. However, it was noted that individuals with a very early onset of SUD, the experience of adverse events was not a significant contributor. One may speculate that this subset of individuals may have stronger genetic underpinnings, and in that respect were more similar to individuals with schizophrenia and BPAD.

Gender differences in the prevalence of ACEs were also noted in our sample, with affected men reporting a greater number of ACEs than affected women. Gender differences in ACEs have been previously described with women more often reporting sexual abuse and household violence and men reporting physical abuse (Edwards et al., 2003). Socio-cultural factors may also explain some of the differences seen in our sample, as it is known that women in the Indian context often do not reveal traumatic experiences owing to stigma and may instead express the same by means of somatic symptoms, which is a cultural idiom of distress (Desai and Chaturvedi, 2017). A study on a community sample also found that males who underwent ACEs were more likely to develop antisocial behaviours, including substance use, in adulthood as compared to females (Schilling et al., 2007).

Interestingly, the association between age of onset and ACE scores in our sample was also seen in men and but not in women. The age of onset of illness, overall, in our sample was also earlier for males than females. This gender specific association of adversity and age of onset could be related to the fact that males with genetic loading for externalising disorders are known to present with more behavioural deviance, and therefore more chastisement by family and alienation from peers, as compared to females who are more likely to develop internalising disorders (Cameron et al., 2017). This may explain an earlier age of onset of the disorders in males in our sample.

The ACE scores of affected individuals in our sample drawn from multiplex families is similar to the ACE scores reported by affected individuals in previous literature (Anda et al., 1999; Chapman et al., 2004; Mersky et al., 2013; Whitfield et al., 2005; Kiburi et al., 2018). This indicates that affected individuals from multiplex families perhaps experience similar levels of adversity, when compared to affected individuals from the general population. The role of ACEs, as an “extra hit” that influences the age of onset in individuals from multiplex families, suggests interplay between environmental adversity during development, and pre-existing vulnerability, that may be mediated by gene-environment interactions. The pathways by which ACEs influence the development of disease phenotypes; including epigenetics, immune-inflammatory, and neuroendocrine mechanisms need further exploration. This could have potential implications for mental health promotion in individuals from multiplex families.

A primary limitation of the study is that the data on ACE recollection is retrospective, and thus may have introduced recall bias. There is also a lack of information in a cultural context as the ACE-IQ scale, while intended for international application, does not address critical dimensions of adversity (such as poverty and food deprivation) that may be more country specific. Additionally, the findings from multiplex families may not be generalizable to individuals without a family history of psychiatric illness.

The findings of the present study provide an understanding of the profile of childhood adversity in individuals from multiplex families. They highlight how exposure to childhood adversity is associated with a younger age of occurrence of a psychiatric condition in these individuals. Future analysis in our longitudinal cohort is expected to identify mechanisms underlying this relationship, specifically in individuals at-risk for developing mental illness.

## Acknowledgements

The authors are grateful to all the patients, and their family members who participated in the study.

## Ethical standards

The authors assert that all procedures contributing to this work comply with the ethical standards laid down by the National Institute of Mental Health and Neuro Sciences (NIMHANS) institutional ethics committee who have approved the study protocol and with the Helsinki Declaration of 1975, as revised in 2008. Written informed consent was obtained from all the participants in the study.

## Data Availability Statement

The data that support the findings of this study are available from the corresponding author upon reasonable request.

---

**Supplementary Table 1:**
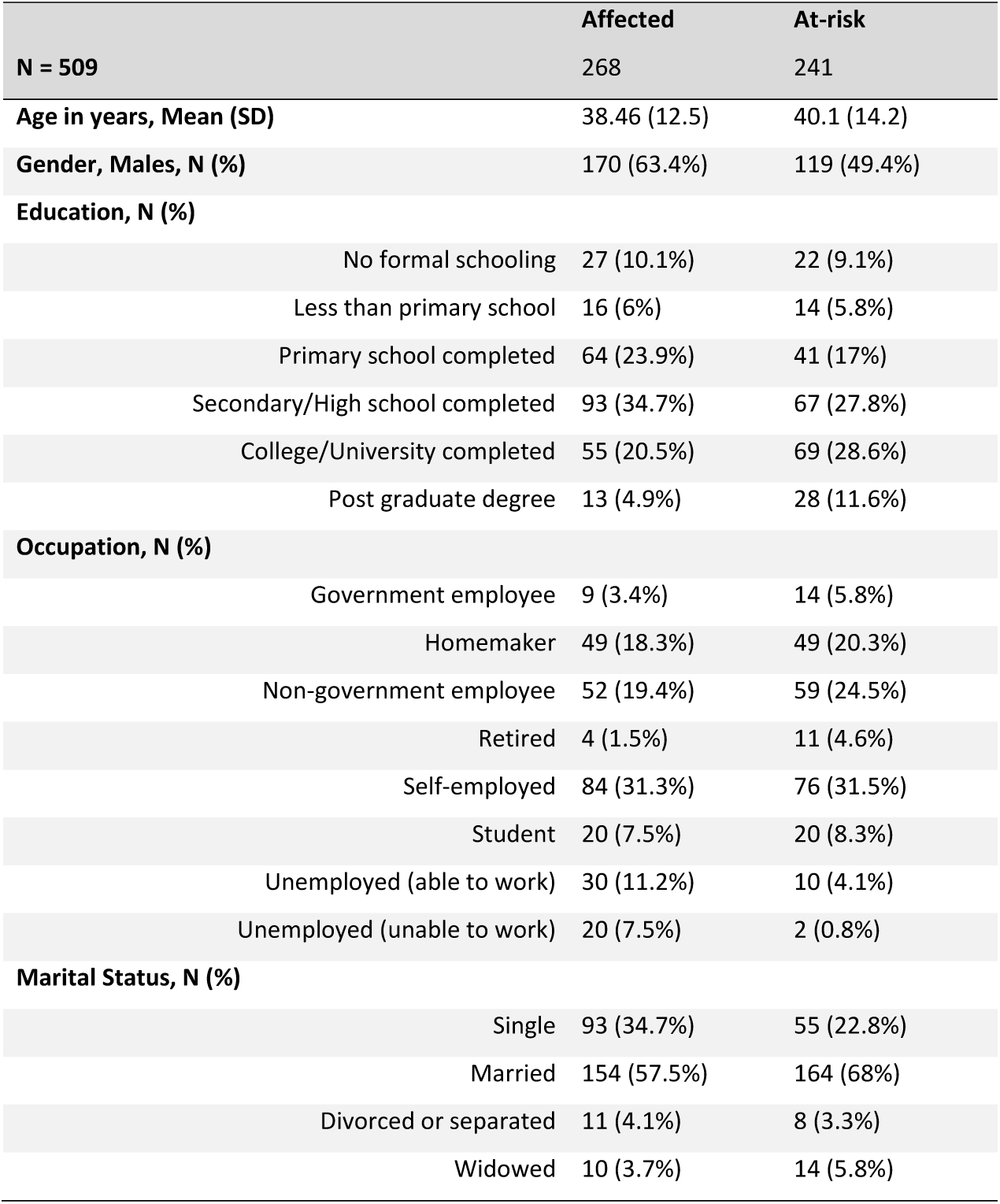
Demographic Profile of the sample.

**Supplementary Table 2:**
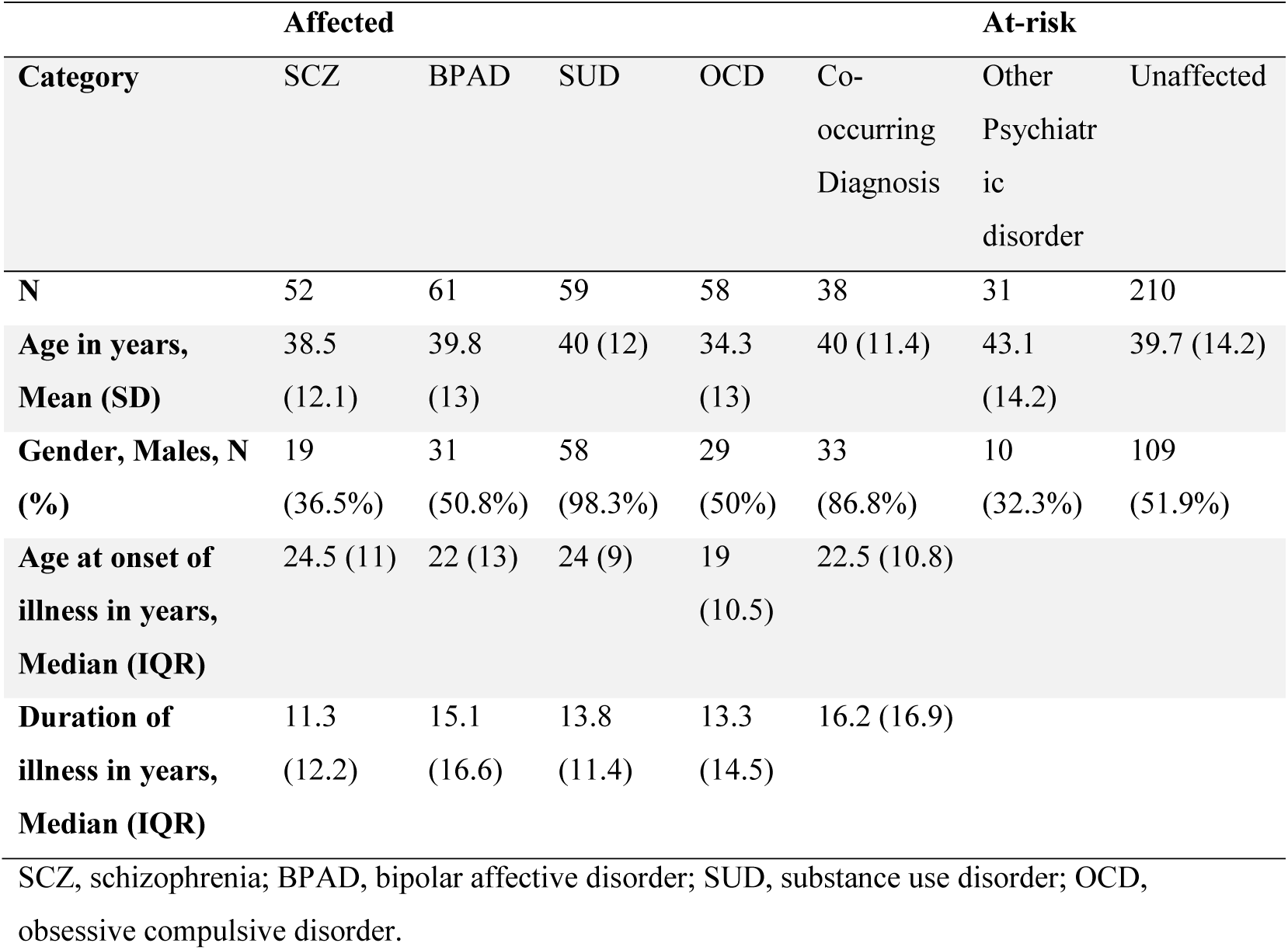
Clinical Profile of the sample.

